# Linking the association between circRNAs and Alzheimer’s disease progression by multi-tissue circular RNA characterization

**DOI:** 10.1101/2019.12.31.892026

**Authors:** IJu Lo, Jamie Hill, Bjarni J. Vilhjálmsson, Jørgen Kjems

## Abstract

Alzheimer’s Disease (AD) has devastating consequences for patients during its slow, progressive course. It is important to understand the pathology of AD onset. Recently, circular RNAs (circRNAs) have been found to participate in many human diseases including cancers and neurodegenerative conditions. In this study, we mined the published dataset on the AMP-AD Knowledge Portal from the Mount Sinai Brain Bank (MSBB) to describe the circRNA profiles at different AD stage in brain samples from four AD patients brain regions, anterior prefrontal cortex, superior temporal lobe, parahippocampal gyrus, and inferior frontal gyrus. We found in total 147 circRNAs to be differentially expressed (DE) during AD progression in the four regions. We also characterized the mRNA-circRNA co-expression network and annotated the potential function of circRNAs based on the co-expressed modules. Based on our results, we propose that parahippocampal gyrus is the most circRNA-regulated region during the AD progression. The strongest negatively AD stage-correlated module in parahippocampal gyrus were enriched in cognitive disability and pathological-associated pathways such as synapse organization and regulation of membrane potential. Finally, the regression model based on the expression pattern of DE circRNAs in the module could help to distinguish the disease severity of patients, further supported the importance of circRNAs in AD pathology. In conclusion, our finding indicates that circRNAs in parahippocampal gyrus are possible regulators of AD progression and potentially be a therapeutic target or of AD.

## Introduction

The global increase in the occurrence of Alzheimer’s disease stresses the urgent need to develop effective medical strategies to help patients suffering from this menacing neurodegenerative disease. Alzheimer’s disease is the most common age-related dementia and it is well known for its slow-progressing and incurable deterioration. The disease manifests as impaired cognitive function caused by neuronal necrosis in specific brain regions(Hyman, Van Hoesen, Damasio, & Barnes, 1984), and eventually leads to death due to brain dysfunction(Pagani et al., 2017). Although the exact pathology is still debated, it is hypothesized that the continuous neuron loss and brain structure dystrophy is induced by the accumulation of Amyloid ß (A ß) protein and cell inflammation(Fang, Sun, Zhou, Qiu, & Peng, 2018), which leads to memory loss and cognitive decline. Appropriate treatment during disease progression could help slow down the disease and it is therefore important to understand the biological and physiological changes that occur during the course of the disease.

Since the discovery of noncoding RNAs (ncRNAs), increasing evidence suggests that ncRNAs play an important role in disease pathology and therefore could serve as both biomarkers and targets for treatment in several human disorders. Recently, ncRNA-related therapeutics targeting cancers, diabetes, and hepatitis C(Rupaimoole & Slack, 2017) have entered the clinical trials. As a new member of the ncRNAs, circular RNAs (circRNA) have caught attention given their abundance in human(Maass et al., 2017) and abnormal behavior in disease states(Han, Chao, & Yao, 2018). CircRNAs were long considered to be by-products of gene expression, however, increasing reports of circRNA dysregulation in cancers(Chen, Shi, Zhang, & Sun, 2017), neurodegenerative diseases(Zhao, Alexandrov, Jaber, & Lukiw, 2016), diabetes(Yonghao Gu et al., 2017) and cardiovascular diseases(Holdt et al., 2016) in the past few years suggest that circRNAs could play an essential role in cell homeostasis and development of diseases(Kristensen et al., 2019).

CircRNAs have covalently linked termini created by a noncanonical back-splicing event and therefore naturally lack free 5’ and 3’ ends. This closed nature of circRNAs provides resistance to exonucleases, thereby increasing their half-life relative to linear transcripts(Enuka et al., 2016), making circRNAs good potential targets for clinical use. Studies on circRNAs as biomarkers have so far mainly been focused on cancers and, to a lesser extent, neurodegenerative disorders. However, there is evidence suggesting the importance of circRNAs in the brain and neural system. The brain is the organ that shows the highest accumulation of circRNAs(You et al., 2015), and many brain-expressed circRNAs are produced from genes with synaptic function in human(You et al., 2015). In addition, expression variation during aging(Gruner, Cortes-Lopez, Cooper, Bauer, & Miura, 2016) has been reported in mammal’s brain. Recently, research demonstrated the correlation between circRNA expression and the pathological features of AD(Dube et al., 2019).

In this study, we set out to investigate how circRNAs correlate with AD progression in different brain regions. We profiled circRNA expression in AD patients using a public RNA sequencing dataset provided by the AMP-AD Knowledge Portal. The dataset was prepared with brain samples from AD patients from Mount Sinai/JJ Peters VA Medical Center Brain Bank (MSBB)(Wang et al., 2018). The dataset contained the Ribo-Zero-treated RNAseq libraries of four brain regions: anterior prefrontal cortex (aPFC), responsible for memory retrieval(Ranganath, Heller, & Wilding, 2007) and prospective memory(Costa et al., 2013), superior temporal lobe (STL) and inferior frontal gyrus (IFG), both involved in language comprehension(Grossman et al., 2002), and parahippocampal gyrus (PHG), essential for episodic memory formation and contextual association(M. Li, Lu, & Zhong, 2016)), from post mortem brain samples at different dementia stages. The dementia stage was determined using the clinical dementia rating (CDR) score(Morris, 1997) provided by the original group. The score is a numeric scale that ranges from zero (normal cognitive ability) to five (terminal stage of dementia) to grade the severity of AD. We identified and compared circRNAs in four human brain regions at different CDR stages and found that PHG was the most influenced region based on the differential expression (DE) analysis and co-expression network results. Functional enrichment analysis of circRNA-mRNA co-expression modules revealed that modules highly associated with disease progression involved pathways including synapse and membrane potential regulation, implying that circRNAs may participate in these pathways. Based on these observations, regression models were built on selected promising circRNA candidates and showed the ability to distinguish between different levels of AD severity, further supporting the importance of PHG circRNAs in AD.

## Results

### Brain region-specific circRNA profiling and changes during disease progression

In order to get an overview of circRNA expression in AD, we first profiled the circRNAs in each brain region. The number of circRNAs identified in the four brain regions ranged from about 58,000 to 72,000, of which 3500 to 5200 circRNAs remained after removing lowly expressed circRNAs in each region (Table 1). About 19% of the identified circRNAs (Supplementary Fig.1) were found in all brain regions, and the percentage of common circRNAs increased to nearly two-thirds among the highly expressed circRNAs (Fig. 1A). In addition, a three-dimensional PCA showed region-specific clustering of the highly expressed circRNAs (Fig. 1B), indicating that both the existence and expression level of circRNAs differed between regions. Among the four regions, IFG showed the highest abundance of circRNAs and clustered separately from the other three regions, suggesting that IFG had a unique expression profile.

**Table 1.**
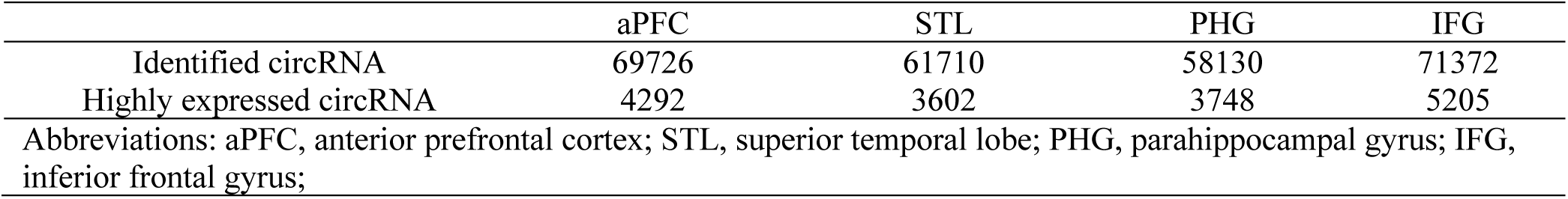
The identified and highly expressed circRNAs in each brain region.

**Figure 1.**
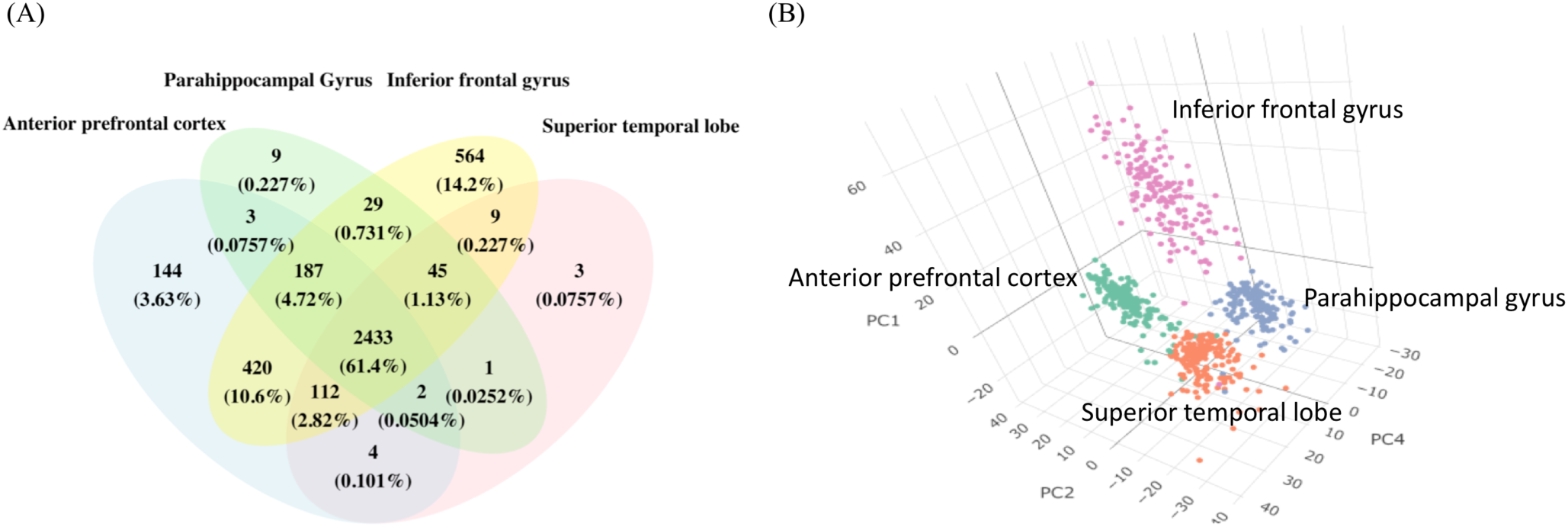
The highly expressed circRNA in four brain regions. (A) Venn diagrams of highly expressed circRNAs in each brain region. The shared and distinct circRNAs are shown as the number and percentage. (B)The 3D PCA of highly expressed circRNAs exists in at least 1 brain regions, the axes show the first, second and fourth principal components.

We then performed differential expression analysis to observe the relationship between circRNA expression and AD progression. We found that the majority of circRNA decreased with increased CDR stage in all four brain regions, irrespective of circRNA expression level and differential enrichment (Supplementary Table 1). A total of 147 circRNAs was significantly DE when the CDR went from zero (no dementia) to five (terminal stage of dementia) at FDR <= 0.05 in all four brain regions (Fig. 2), but the distribution of the 147 circRNAs was highly specific. Only four of the 147 differentially expressed circRNAs (2.7%) are shared across the four brain regions, and the number of region-specific differentially expressed circRNAs are 19 (12.9%), 9 (6.12%), 54 (23.8%) and 35 (36.7%) in aPFC, STL, PHG, and IFG, respectively (Supplementary Fig. 2). In each brain region, 35,21,77 and 61 circRNAs were differentially expressed. Among these significantly differentially expressed circRNAs, 12,7,33,21 were up-regulated, and 23,14,44,40 circRNAs were downregulated with the progression of dementia in aPFC, STL, PHG, and IFG, respectively. Although the profiling data showed that IFG had the highest circRNA expression, PHG was the region with the most significantly changed circRNAs during the AD progression.

**Figure 2.**
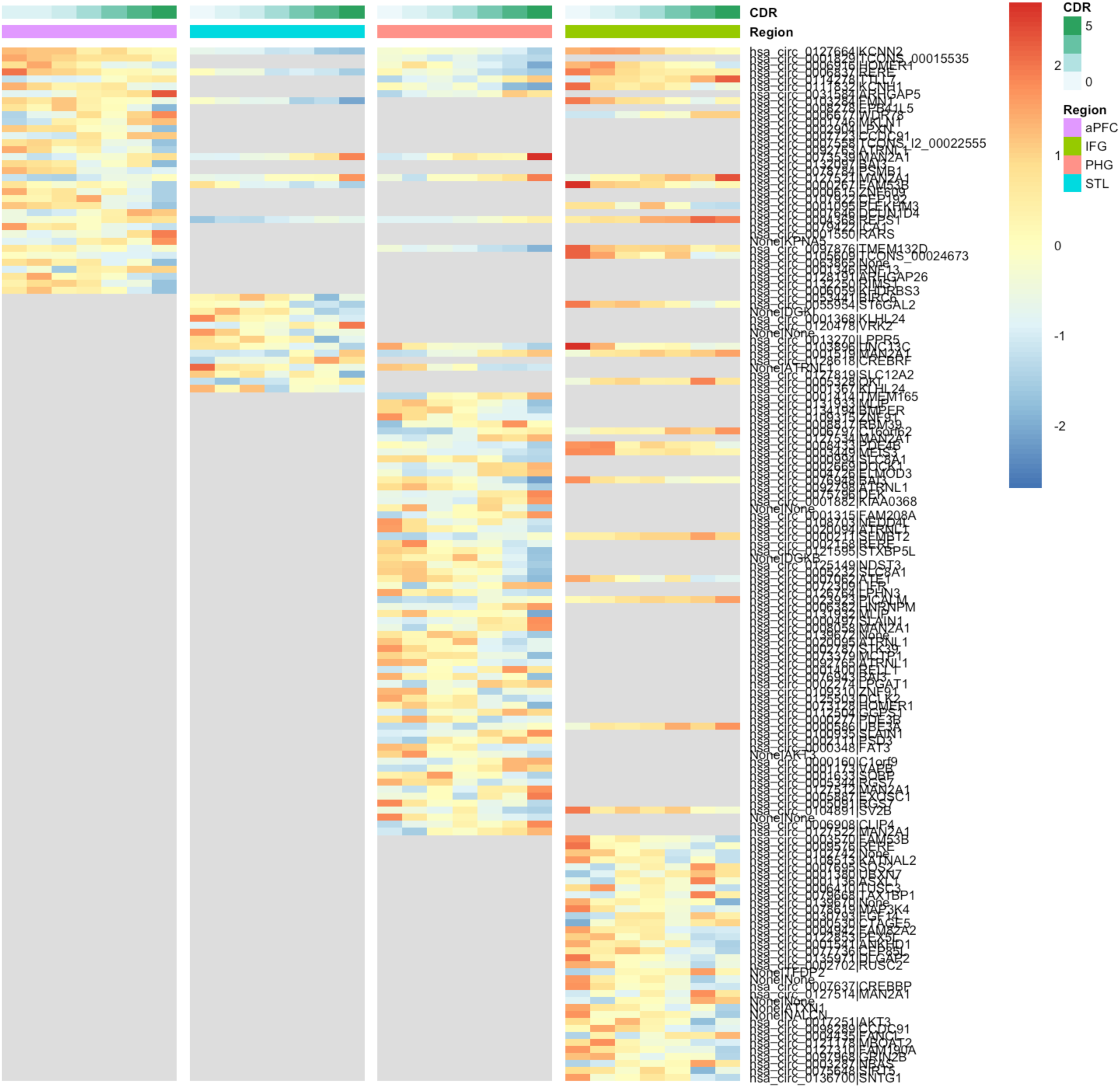
The heatmap of significantly DE circRNAs in brain regions. The color legends on top of the heatmap show the CDR stages and the brain region. The grey area in the plot means the circRNA is not significantly DE in that region.

### circRNA-mRNA co-expression network analysis and functional annotation

Given the global deregulation in circRNA expression during AD progression we speculated that a correlated dysregulation of circRNAs and mRNAs during cognitive decline could suggest a functional role in shared pathways. We therefore constructed a circRNA-mRNA co-expression network to capture the relationship between circRNAs and mRNAs. Moreover, the co-expression network aided in the functional enrichment test for circRNAs. Despite numerous reports suggesting the function of circRNAs, the experimentally validated functional annotation of most circRNAs is still far from completion. Therefore, considering that the mRNAs and circRNAs in the same module could be involved in similar mechanisms during disease progression, we assumed that the mRNAs in a module could be used to estimate circRNA function.

Fig 3A shows the correlation heatmap of circRNA-mRNA co-expressed modules and CDR in PHG, including sex and AOD as they are strong influence factors of AD. The topological matrix of the same region is shown in Fig 3B. The co-expressed modules were assigned to different colors to visualize the topological overlap, but these do not represent any functional information. (The correlation heatmap and the topological matrix of the other three regions are in Supplementary Figure 3, and the composition of modules in Supplementary Table 2∼5). All modules in aPFC and STL were weakly associated with CDR, consistent with a previous study showing that these two regions were the least important of the four brain regions in dementia(Wang et al., 2016). Surprisingly, although IFG was shown to be an important region for AD in the previous study, and it was the region with the highest circRNA expression in our study, the association between the IFG transcriptome and the CDR score was nearly as weak as for aPFC and STL. The maximum absolute correlation coefficients of the three regions were 0.32, compared to 0.51 for the PHG. This, however, agreed with the observation that PHG had more significantly DE circRNAs than all the other 3 regions, suggesting that circRNAs in PHG were most associated with AD progression. We therefore focused on the functional role of circRNAs in this specific region. In PHG, the module with highest correlation coefficient was ‘violet’ (r = 0.49 and *P* value = 8^10-10) and the lowest was ‘brown’ (r = - 0.51 and *P* value = 2^10-10).

**Figure 3.**
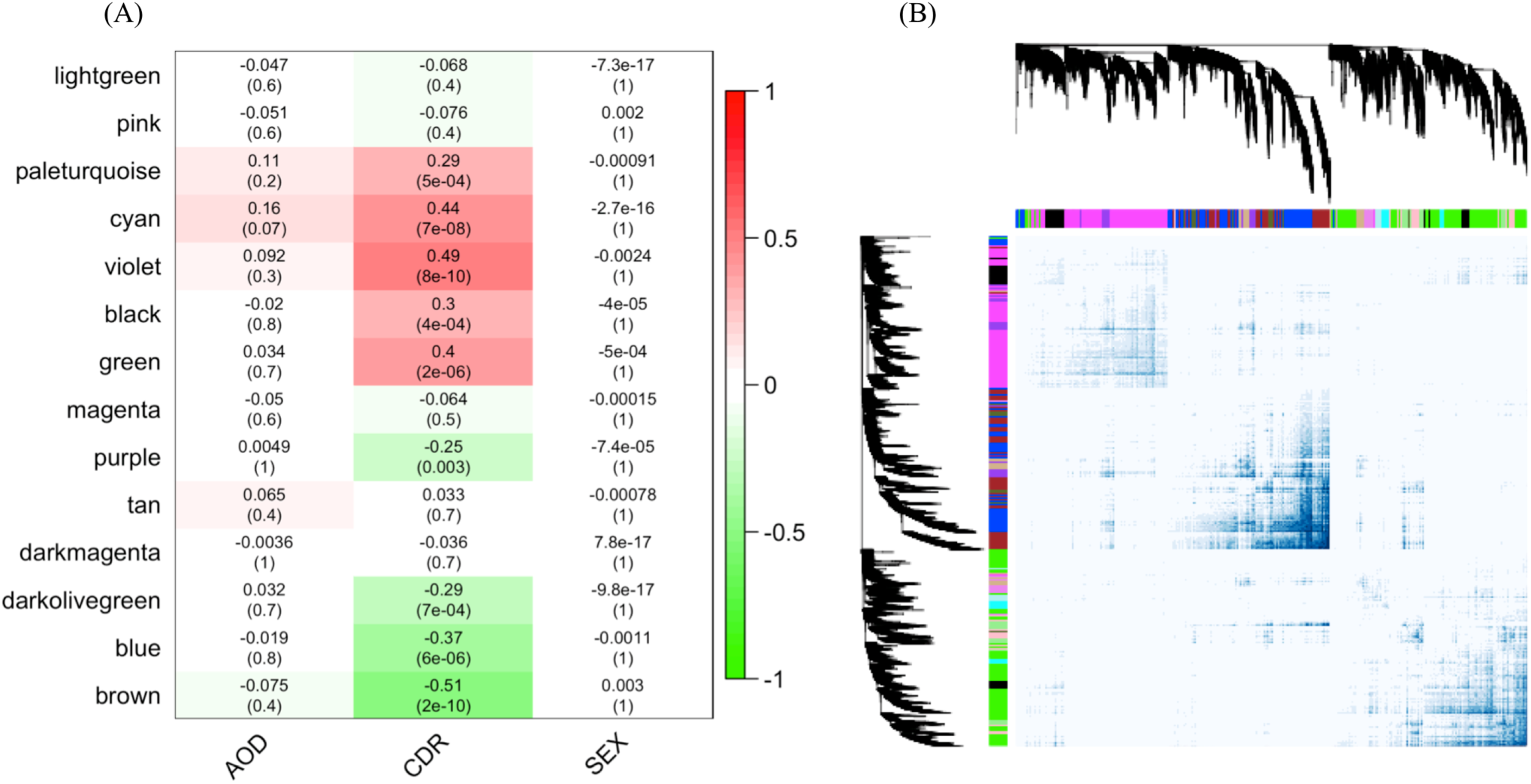
The circRNA-mRNA co-expressed modules in parahippocampal gyrus. (A) represents the correlation of circRNA-mRNA co-expressed modules clustered by WGCNA in the parahippocampal gyrus and three traits, CDR, AOD, and SEX. The numbers in the heatmaps are the correlation coefficients and the *P* value in the parentheses. (B) shows the topological overlap of the co-expressed modules. Each column and row is a transcript and the color bands on the top and left represent the modules. The deeper colored regions in the network matrix correspond to more topological overlap (i.e. higher intra-module similarity of the members). The dendrograms on top and left show the clustering of the transcripts.

For each module, the host genes of mRNAs and circRNAs that significantly changed with the CDR were then extracted for GO and DO analysis. The brown module was enriched in important AD pathological mechanisms such as membrane potential regulation and synaptic function, and advanced cognitive functions including behavior, learning or memory and cognition(Supplementary Figure 4A). The DO enrichment test also showed that the brown module was significantly related to mental disorders such as attention deficit hyperactivity disorder, autism, depression, and epilepsy (Supplementary Figure 4B). Interestingly, weight change and, specifically, weight gain were enriched DO terms in our analysis. Although it is not a feature most commonly linked to Alzheimer’s disease, several studies have reported that before and during Alzheimer’s disease progression, some patients experience unusual and dramatic weight change (Yian Gu et al., 2014). The networks in Fig 4A and 4B represented the strongest interaction between circRNAs and mRNAs in the brown module, color-coded by the ontology enrichment terms. Comparing to the brown module, the violet module was less functional enriched, the only significant biological pathway enriched term was epithelial cell differentiation, and circRNA-mRNA correlation in the network was much weaker than in the brown module (data was not shown).

**Figure 4.**
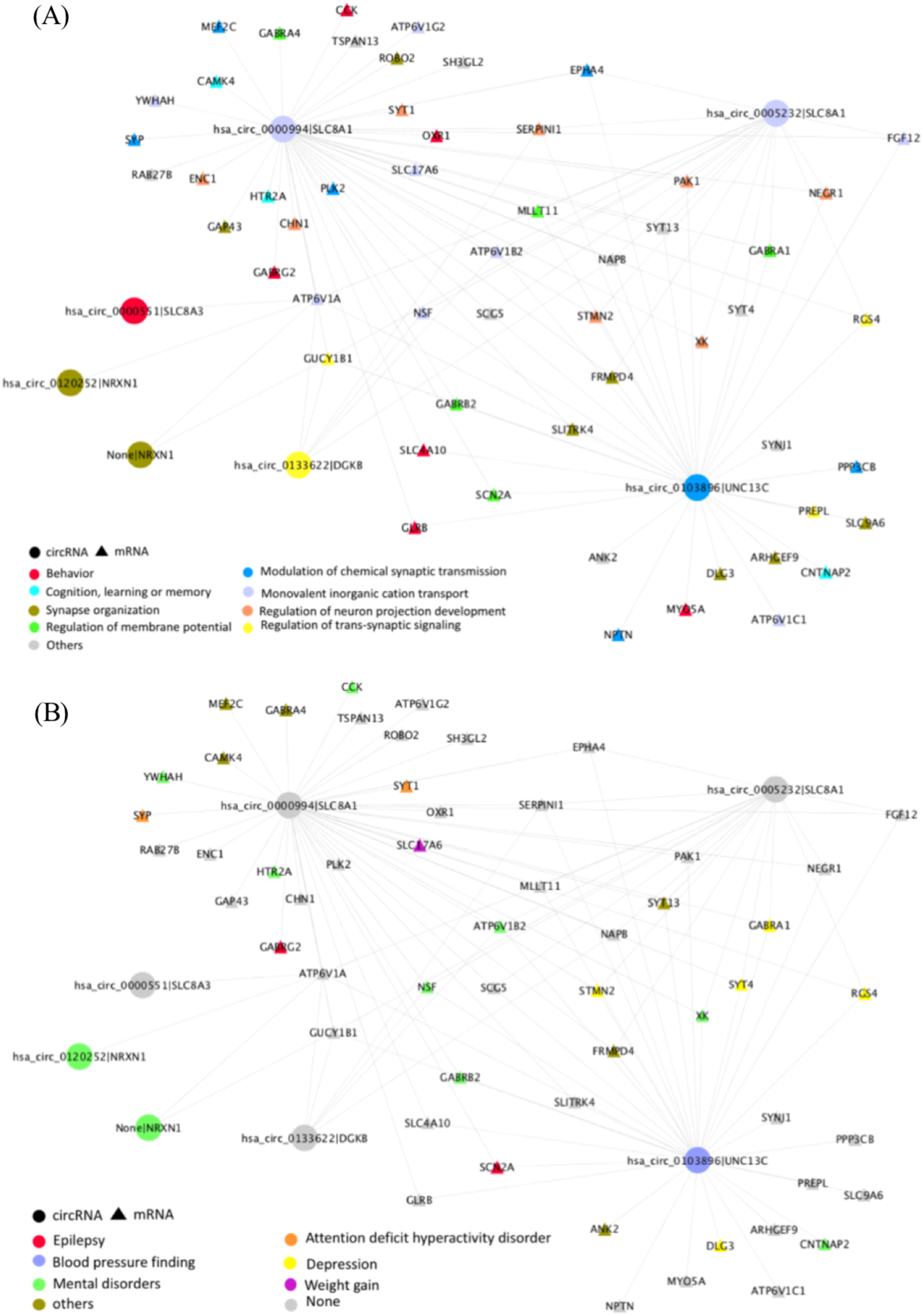
The circRNA-mRNA co-expressed network of the brown module in Parahippocampal gyrus. The same network color code by either (A) biological pathway (B) disease ontology enrichment term. The circle represents the circRNAs and the triangle represents the genes. Only the circRNAs and mRNAs which log2 fold change >= 0.2 & FDR <= 0.05, the edge weight >= 0.2 are used, and the mRNA-mRNA interactions are excluded to provide a brief overview.

### Association between circRNAs and Alsheimer’s disease progression

Given that circRNAs in the PHG brown module are related to disease progression, we selected 13 circRNAs in the PHG brown module (see methods for specific criteria) as promising candidates involved in the pathological process. We grouped individuals into 3 CDR groups by severity and plotted the expression differences for the 13 circRNAs in Supplementary Figure 5. We then used a multinomial regression model to quantify how well the 13 circRNAs predicted any of the three AD severity groups (model and analysis details are described in the methods section). The result is shown in Supplementary figure 6. To further examine the association between circRNAs and AD progression, we selected a subset of five circRNAs that each had an area under the ROC curve (AUC) >= 0.7 in Supplementary Figure 6 and found their joint predictive value for AD to be significant (p = 0.035). The ROC curve for this model is shown in Fig. 5. The average AUC of 5-circRNA model is 0.79, comparing to the 0.69 average AUC of the base model. Moreover, in each group comparison, the 5-circRNA model showed improvement from the base model. For healthy/MCI, MCI/AD, and healthy/AD comparison, the 5-circRNA model had AUC 0.64, 0.76, and 0.96, while the base model had AUC 0.57, 0.62 and 0.89, respectively. The better performance of the 5-circRNA models suggesting that these circRNAs could be important for AD progression, and the potential to help to improve the diagnostics with the better understanding of the biological mechanisms these circRNAs involved in.

**Figure 5.**
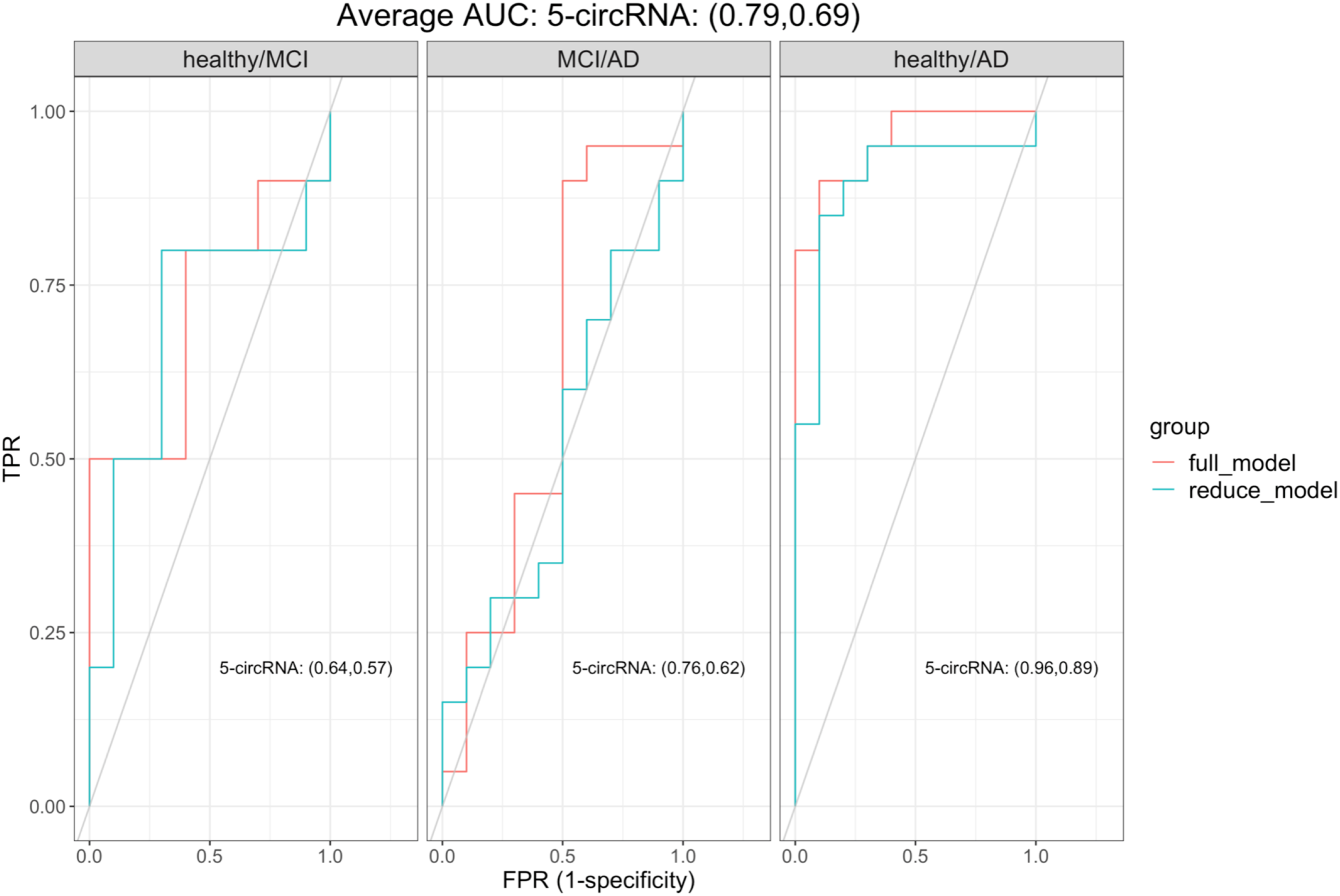
The ROC of the multivariate regression model of circRNA candidates. The 5-circRNA model constructed from five circRNAs with AUC >= 0.7 in Supplementary Fig 6. The average AUC of three comparisons annotated at the top of the figure, and the AUC of each comparison annotated at the bottom right corner (the former number in the parentheses were AUC of the full model and the latter were AUC of the reduced model). The color indicates the used model.

## Discussion

In this study, we identified a global dysregulation of circRNAs in four brain regions during AD progression, especially a decreased circRNA expression in diseased brains compared to the general increase of circRNAs seen during normal aging(Gruner et al., 2016), suggesting that circRNAs may be beneficial for healthy aging.

Comparing all four regions studied, we found that although IFG was the region with the most enriched circRNA expression, PHG was the region in which circRNAs were most related to the cognitive decline in AD, and the circRNAs in PHG were enriched in AD pathological pathway. Considering that PHG is mainly responsible for advanced cognitive abilities (e.g, associative learning), and that many reports have indicated that PHG atrophy could be an early symptom of AD patients(Echávarri et al., 2011; Krumm et al., 2016), the abnormal change of circRNAs implies a functional association of circRNAs in PHG and AD progression. We also built a regression model to test the association between circRNAs and AD. Due to the limited number of available independent datasets, we used 70% of the samples in our dataset for model training and 30% for the performance test, while all samples were used for DE and co-expression network analysis. The test data for the regression model was therefore not fully independent, and further validation is necessary to confirm the results. Nevertheless, the regression model showed that the selected circRNAs in PHG improved the classification of patients at different AD stages. Our results support that the selected circRNAs might be involved in the AD progressive pathology and could be shortlisted candidates for experimental analyses.

Finally, considering the high stability of circRNAs and that circRNAs have been found to circulate in the bloodstream encapsulated in exosomes(Xu, Guo, Li, & Yu, 2015), it is possible that circRNA could be used for future diagnostic approaches for AD based on blood samples. Although it is necessary to further confirm the robustness of the circRNA candidates described here and determine if they can be detected in exosomes or biofluids, our research describes the AD-associated circRNA changes and pointed out the potential importance of PHG circRNAs during the AD disease process. The results could also provide biological insights to narrow down candidates for further studying circRNAs and potential AD therapeutic targets.

## Methods

### Dataset information

The clinical information, RNA, and QC tables were downloaded from the AMPAD Knowledge Portal (https://www.synapse.org/#!Synapse:syn3159438) in June 2018. As part of the MSBB study, there were 364 individuals at different AD stages in the cohort. Ribo-Zero-treated 100 bp single-end RNA sequencing libraries were available for one to four of the below brain regions for each patient: aPFC, STL, PHG, and IFG. The detailed tissue processing and data preparation procedures have been described(Wang et al., 2018). Not all subjects had RNA-Seq libraries available for all four brain regions, and not all the libraries passed the quality control performed by the original group. To avoid low-quality samples, individuals who were not annotated as “Okay” in the QC table provided by the original group, samples where the RNA sequencing had a RIN score lower than 4, or samples where the rRNA rate was higher than 5% were removed. Samples from non-European individuals were also removed. Finally, of 223 individuals, there were 187,166,138 and 158 RNA sequencing datasets available in four regions, respectively. The demographic summary of the dataset is provided in Supplementary Table 6.

### CircRNA quantification

To quantify circRNA expression in the AD brain regions, the BAM files and the unmatched FastQ files (containing reads that failed to map to the reference genome) of the same individual were downloaded, converted and merged back to the FastQ file containing all sequenced reads with Picard(v2.7.1)(“Picard Tools - By Broad Institute,” n.d.) and then re-mapped to the human reference genome hg19 with the Burrows-Wheeler Aligner (BWA-MEM, v0.7.15)(H. Li, 2013). CircRNA detection tool CIRI2(v2.0.6)(Gao, Zhang, & Zhao, 2018) was used to identify and quantify the circRNAs. In order to construct co-expression networks, we also downloaded the mRNA expression tables from the AMPAD Knowledge Portal. Further data cleaning and analyses were done using R.

The R package org.Hs.eg.db(M. Carlson, 2018), TxDb.Hsapiens.UCSC.hg19.knownGene(M. and B. P. M. Carlson, 2015), Ensembl genomic information from the BioMart portal(www.biomart.org)(Smedley et al., 2015), and information of known circRNAs from circBase(http://circbase.org/)(Glažar, Papavasileiou, & Rajewsky, 2014) were used for circRNA and mRNA annotation. After annotation, transcripts that passed both of the following criteria: (1) transcripts with zero reads in less than 25% of individuals (2) average circRNA expression higher than two reads (mRNA higher than ten reads) in any CDR group were considered as highly expressed (The full list of circRNAs passed the criteria is provided as Supplementary Material Supplementary_Table_S7.xlsx). The highly expressed circRNAs of all samples were then used for principal component analysis (PCA) to find potential clustering. All the following analyses also used only highly expressed transcripts unless otherwise noted.

### Differential expression analysis

DE analysis of circRNA was performed with the R package DESeq2(v1.18.1)(Love, Huber, & Anders, 2014). The model used CDR score as the dependent variable and included the postmortem interval (PMI), age of death (AOD), sex and batch information as covariates. *P* values were adjusted for multiple testing using the Benjamini-Hochberg (BH) method. Only results with an adjusted *P* value <= 0.05 were considered significant. (The DE result of all circRNAs is provided as Supplementary Material Supplementary_Table_S8.xlsx)

### Co-expression network construction

The R package WGCNA (v1.66)(Langfelder & Horvath, 2008) was used to construct circRNA-mRNA co-expression networks. Before running WGCNA, PCA analysis was performed to check for potential confounder effects in circRNA and mRNA in each brain region, and clustering based on the batch in circRNA and sex in mRNA was detected. Since the sex effect was not the focus in this study, the batch effect in circRNAs and sex effect in mRNAs were removed with the Limma package (v3.38.3)(Ritchie et al., 2015)(Supplementary Fig 7); the network construction then followed the tutorial provided by the WGCNA package. Briefly, using default parameters, WGCNA calculated and picked a soft threshold for each brain region and removed outliers. WGCNA then constructed a signed network based on the expression change of all highly expressed circRNAs and mRNAs and calculated the Pearson correlation between the clustered models and the CDR. To avoid too many small modules, the clustered dendrogram was trimmed to keep modules containing at least 100 transcripts. Network visualization was done by Cytoscape (v3.7.1)(Shannon et al., 2003). Only those circRNAs and mRNAs with a log2-fold change >= 0.2 & FDR <= 0.05, and the edge weight >= 0.2 were used for network visualization; mRNA-mRNA interactions were excluded to provide a simplified overview.

### Functional enrichment analysis

Gene ontology (GO) and disease ontology (DO) enrichment analyses were performed by the R packages clusterProfiler(v3.10.1)(Yu, Wang, Han, & He, 2012) and DOSE(v3.8.2)(Yu, Wang, Yan, & He, 2015). Given the sparsity of functional information for circRNAs, the mRNAs that were significantly changed during cognitive decline (FDR <= 0.05) and in the most positively- or negatively CDR-correlated co-expression modules were also included in the functional enrichment test, while all highly expressed mRNAs and circRNAs were used as the background for the test. The redundant enriched ontology terms were trimmed by the clusterProfiler, and the remaining terms with *P* value <0.05 were selected.

### Regression model construction and the receiver operating characteristics curve (ROC)

We found 13 circRNAs that passed both the following criteria: (1) significantly DE during cognitive decline (FDR <= 0.05 and log2 fold change >= 0.2). (2) member of the most trait-related WGCNA co-expression modules. These 13 circRNAs were analyzed further in a multinomial regression model. All individuals were categorized as three disease stages: healthy (CDR less than one), mild cognitive impairment (MCI, CDR between one and three) and AD (CDR higher than three), which was then used as the outcome in the model. We included the following covariates in the base model: age of death (AOD), sex, batch and the first 3 principal components of circRNA expression. We then examined the relative area under the receiver operating characteristics curve (AUC) for each single circRNA candidate, i.e. the increase in AUC obtained by including the circRNA expression as a covariate in the base model. We employed R package caret(v6.0-84)(Kuhn, 2008) and glmnet(v2.0-18)(Friedman, Hastie, & Tibshirani, 2010) for the analysis. For model training, 10-fold cross-validation performed on 70% of individuals selected by stratified random sampling by the disease stage. The rest of 30% of individuals were used as test data for model performance estimation. The number of samples in each group in training and testing datasets for the regression model are listed in Supplementary Table 9. The five most strongly associated circRNAs among the 13 examined individually in the regression were subsequently selected for further analysis. Using the same base model, we included the five circRNAs as covariates and examined the relative AUC.

## Data access

MSBB: https://www.synapse.org/#!Synapse:syn3159438

## Funding

This project has received funding from the European Union’s Horizon 2020 research and innovation program under the Marie Sklodowska-Curie grant agreement No 721890

The results published here are in whole or in part based on data obtained from the AMP-AD Knowledge Portal (doi:10.7303/syn2580853). These data were generated from postmortem brain tissue collected through the Mount Sinai VA Medical Center Brain Bank and were provided by Dr. Eric Schadt from Mount Sinai School of Medicine.

We thank for the patients and researchers who contributed to generating this high-quality dataset, without them this study would not be possible. We also thank Dr. Anne Færch Nielsen for critically reading of the manuscript, the whole Kjems lab, and QIAGEN Aarhus for constructive comments, discussion, and support.

## Disclosure declaration

The authors declare that there are no conflict of interests.

## Supplementary

**Supplementary Table 1.** The differentially expressed circRNAs in each brain region.

|  | aPFC | STL | PHG | IFG |
| --- | --- | --- | --- | --- |
| Up-regulated DE circRNAs | 1954 | 1550 | 1681 | 2480 |
| Down-regulated DE circRNAs | 2338 | 2052 | 2067 | 2725 |
| Total | 4292 | 3602 | 3748 | 5205 |

Abbreviations: aPFC, anterior prefrontal cortex; STL, superior temporal lobe; PHG, parahippocampal gyrus; IFG, inferior frontal gyrus; DE, differentially expressed;
NOTE. The numbers represent in highly expressed circRNAs, how many circRNAs have positive log2 fold change (up-regulated) or negative log2 fold change (down-regulated) during the CDR stages, no matter it is significantly at FDR =0.05 or not.

**Supplementary Table 2.** The WGCNA module composition of the anterior prefrontal cortex.

| ModuleColor | circRNA | mRNA | Total |
| --- | --- | --- | --- |
| black | 147 | 2323 | 2470 |
| blue | 764 | 6551 | 7315 |
| brown | 946 | 2578 | 3524 |
| cyan | 0 | 185 | 185 |
| green | 0 | 3532 | 3532 |
| greenyellow | 0 | 279 | 279 |
| grey60 | 0 | 116 | 116 |
| lightcyan | 0 | 126 | 126 |
| salmon | 0 | 230 | 230 |
| tan | 1 | 742 | 743 |
| turquoise | 2434 | 6001 | 8435 |

**Supplementary Table 3.**
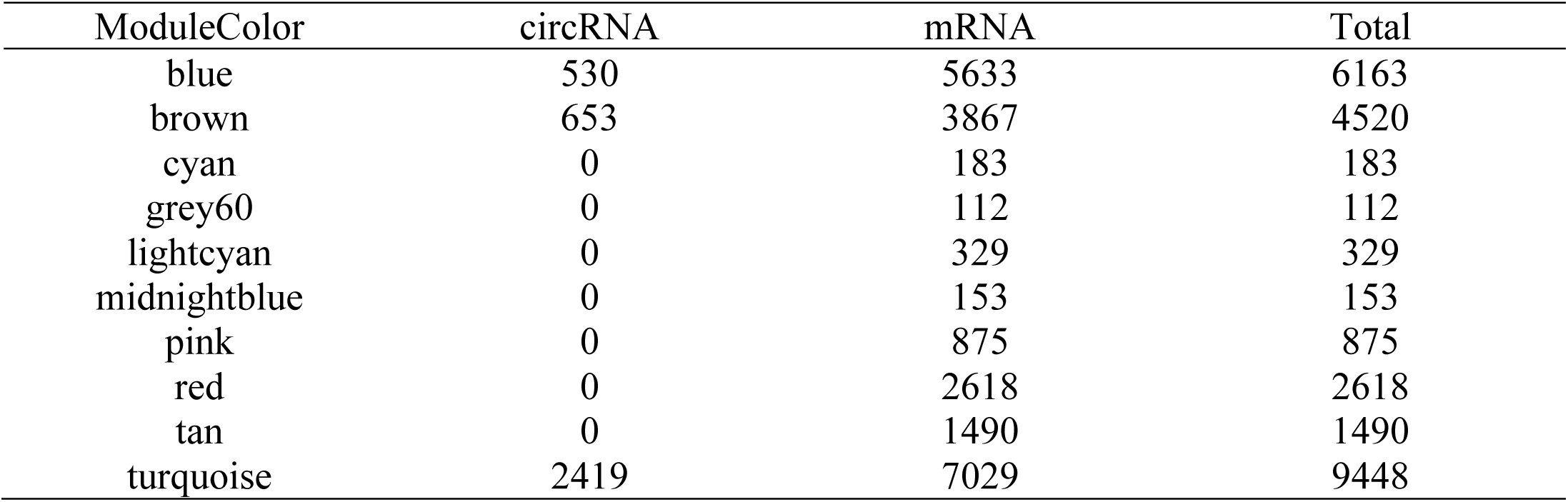
The WGCNA module composition of the superior temporal lobe.

| ModuleColor | circRNA | mRNA | Total |
| --- | --- | --- | --- |
| blue | 530 | 5633 | 6163 |
| brown | 653 | 3867 | 4520 |
| cyan | 0 | 183 | 183 |
| grey60 | 0 | 112 | 112 |
| lightcyan | 0 | 329 | 329 |
| midnightblue | 0 | 153 | 153 |
| pink | 0 | 875 | 875 |
| red | 0 | 2618 | 2618 |
| tan | 0 | 1490 | 1490 |
| turquoise | 2419 | 7029 | 9448 |

**Supplementary Table 4.** The WGCNA module composition of the parahippocampal gyrus.

| ModuleColor | circRNA | mRNA | Total |
| --- | --- | --- | --- |
| black | 11 | 1546 | 1557 |
| blue | 1130 | 2296 | 3426 |
| brown | 1171 | 2409 | 3580 |
| cyan | 0 | 687 | 687 |
| darkmagenta | 0 | 151 | 151 |
| darkolivegreen | 0 | 675 | 675 |
| green | 1174 | 4395 | 5569 |
| lightgreen | 0 | 949 | 949 |
| magenta | 56 | 5498 | 5554 |
| paleturquoise | 22 | 420 | 442 |
| pink | 85 | 1090 | 1175 |
| purple | 5 | 1073 | 1078 |
| tan | 63 | 937 | 1000 |
| violet | 31 | 589 | 620 |

**Supplementary Table 5.** The WGCNA module composition of the inferior frontal gyrus.

| ModuleColor | circRNA | mRNA | Total |
| --- | --- | --- | --- |
| black | 0 | 192 | 192 |
| blue | 2157 | 10004 | 12161 |
| brown | 40 | 1136 | 1176 |
| pink | 0 | 123 | 123 |
| red | 53 | 962 | 1015 |
| turquoise | 2955 | 9489 | 12444 |

**Supplementary Table 6.** Demographic summary of patients used in the study.

|  |  | aPFC | STL | PHG | IFG |
| --- | --- | --- | --- | --- | --- |
| Sex | Male | 66 | 61 | 53 | 55 |
|  | Female | 121 | 105 | 85 | 103 |
| AOD (year) | | 84.18 (± 7.03) | 83.89(± 7.31) | 83.98(± 7.39) | 84.44(± 7.03) |
| CDR<br>(0 = healthy, 5 =<br>terminal dementia) | 0 | 25 | 22 | 19 | 19 |
|  | 0.5 | 22 | 19 | 15 | 19 |
|  | 1 | 18 | 19 | 16 | 18 |
|  | 2 | 31 | 22 | 19 | 25 |
|  | 3 | 40 | 41 | 25 | 35 |
|  | 4 | 24 | 18 | 21 | 21 |
|  | 5 | 27 | 25 | 23 | 21 |
| Total |  | 187 | 166 | 138 | 158 |

Abbreviations: aPFC, anterior prefrontal cortex; STL, superior temporal lobe; PHG, parahippocampal gyrus; IFG, inferior frontal gyrus; AOD, age of death; CDR, clinical dementia rating.
NOTE. According to the HIPAA compliance, individuals whose AOD ≥ 90 have been censored to 90+, and for mean AOD calculation were encoded as 90 years. The numbers in the AOD column are the average, and standard deviation in the parentheses.

**Supplementary Table 9.** Number of individuals used for multivariate logistic regression.

|  | Healthy | MCI | AD |
| --- | --- | --- | --- |
| Training | 24 | 25 | 49 |
| Test | 10 | 10 | 20 |
| Total | 34 | 35 | 69 |

**Supplementary Figure 1.**
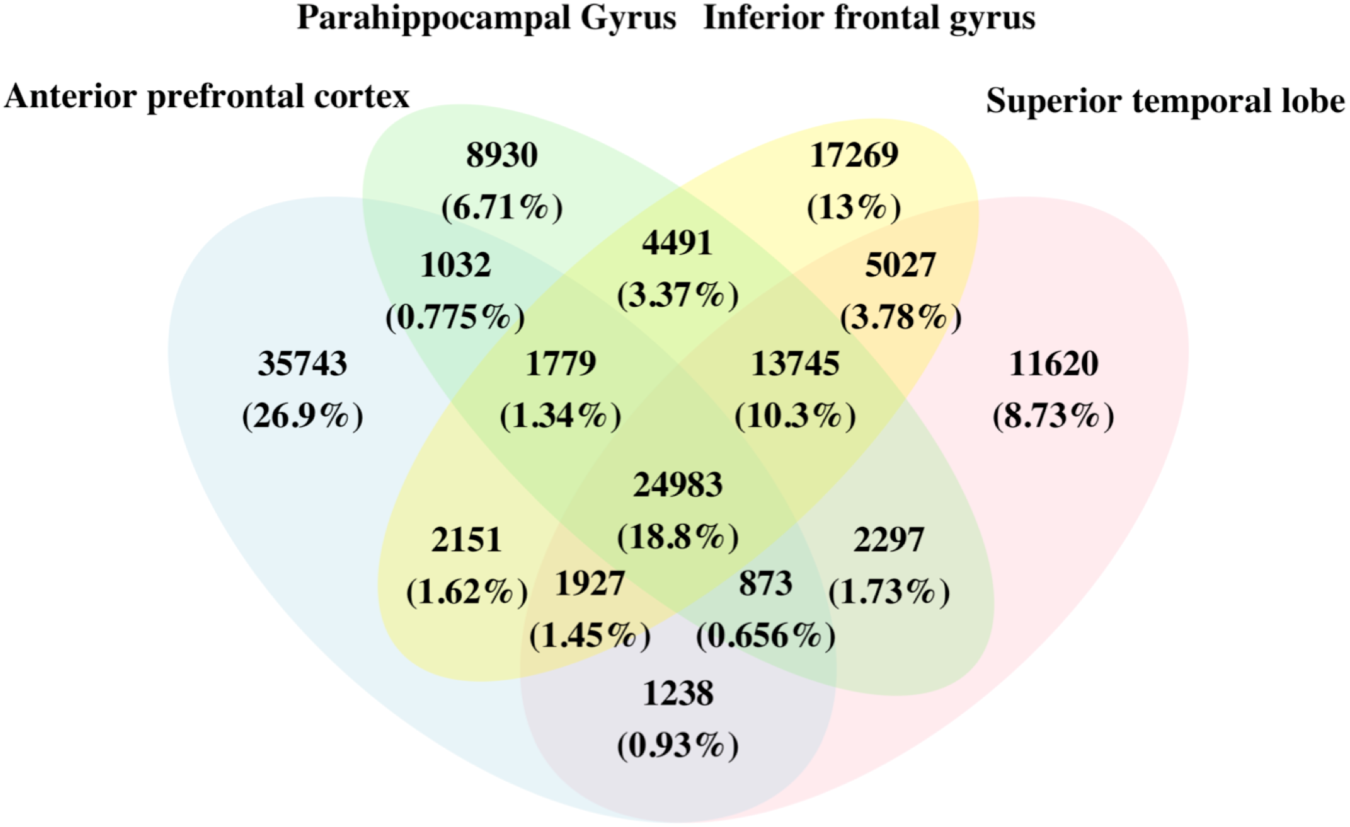
The distribution of identified circRNAs in brain regions. The numbers and the percentages show the distribution of all circRNAs identified by CIRI2 software among four brain regions

**Supplementary Figure 2.**
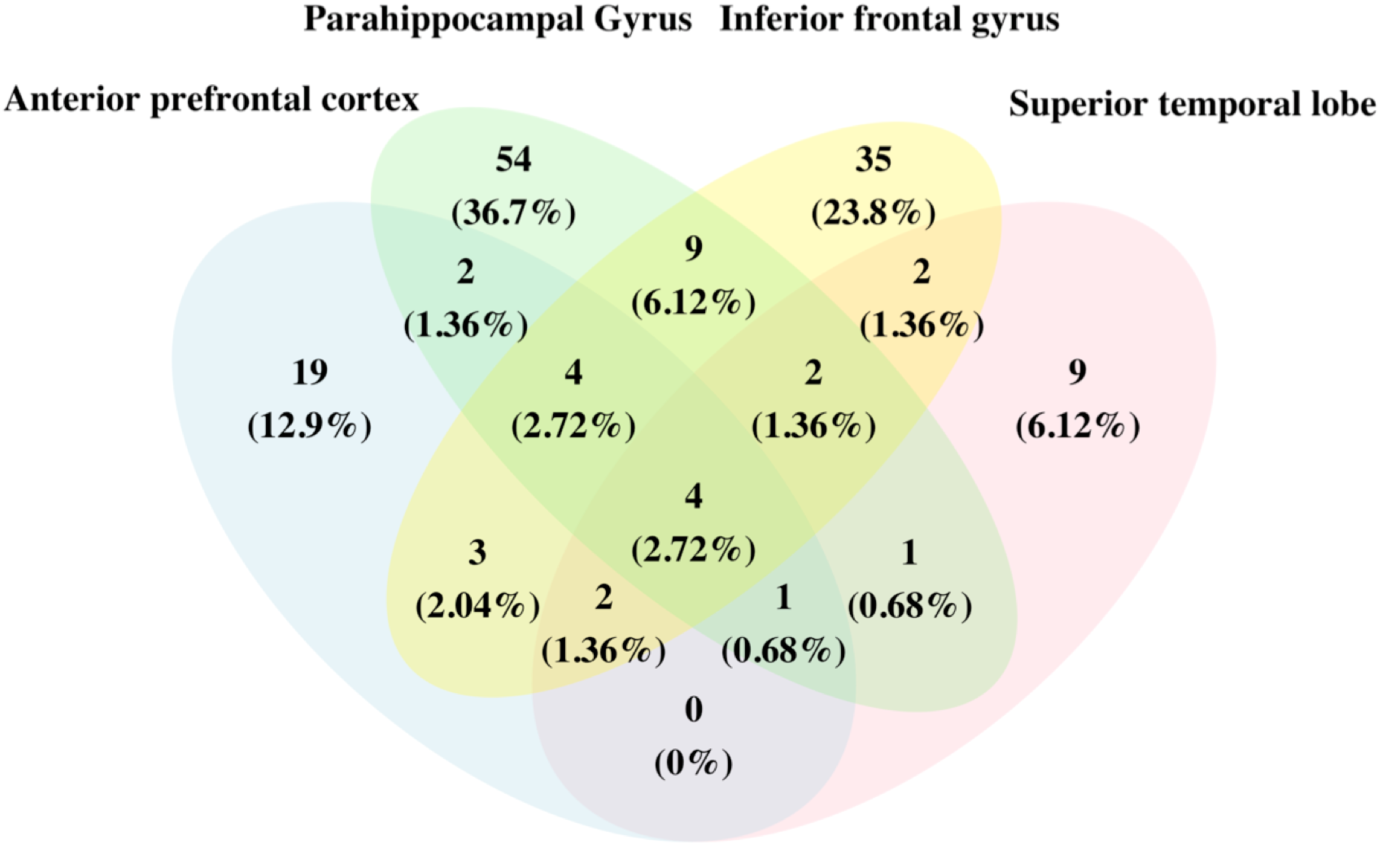
The Venn diagram of significantly differentially expressed (DE) circRNAs. The numbers and the percentages represent how many circRNAs are significant DE at FDR <= 0.05 among four brain regions.

**Supplementary Figure 3.**
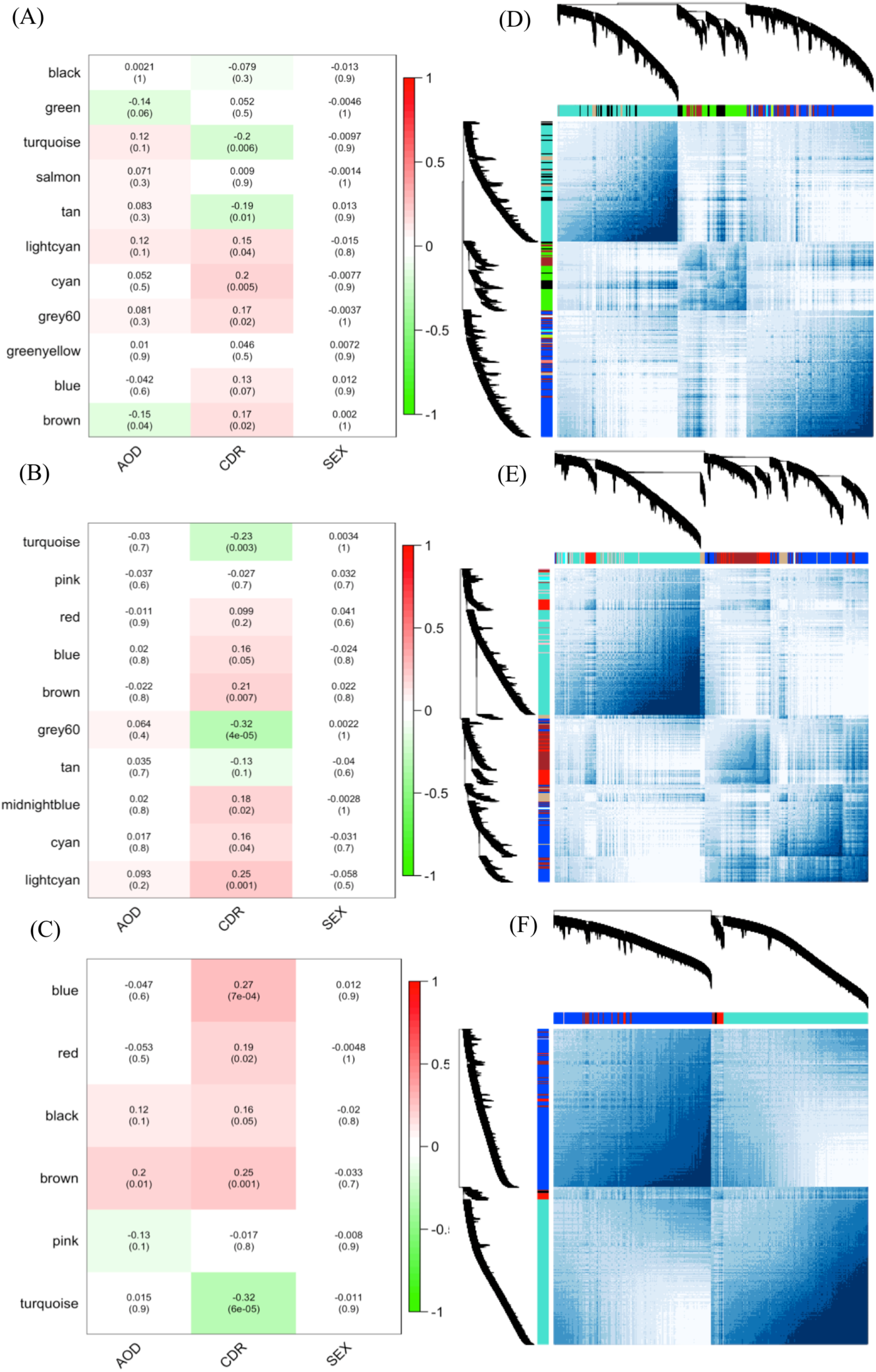
The circRNA-mRNA co-expressed modules in the anterior prefrontal cortex (aPFC), superior temporal lobe (STL) and inferior frontal gyrus (IFG). (A) –(C) represent the correlation of co-expressed modules clustered by WGCNA in the aPFC, STL and IFG to the three traits, CDR, AOD, and SEX. The numbers in the heatmaps are the correlation coefficients and the *P* value in the parentheses. (D) – (F) show the topological overlap of the co-expressed modules in aPFC, STL and IFG, respectively. Each column and row is a transcript and the color bands on the top and left represent the modules. The deeper colored regions in the network matrix correspond to more topological overlap (i.e. higher intra-module similarity of the members). The dendrograms on top and left show the clustering of the transcripts.

**Supplementary Figure 4.**
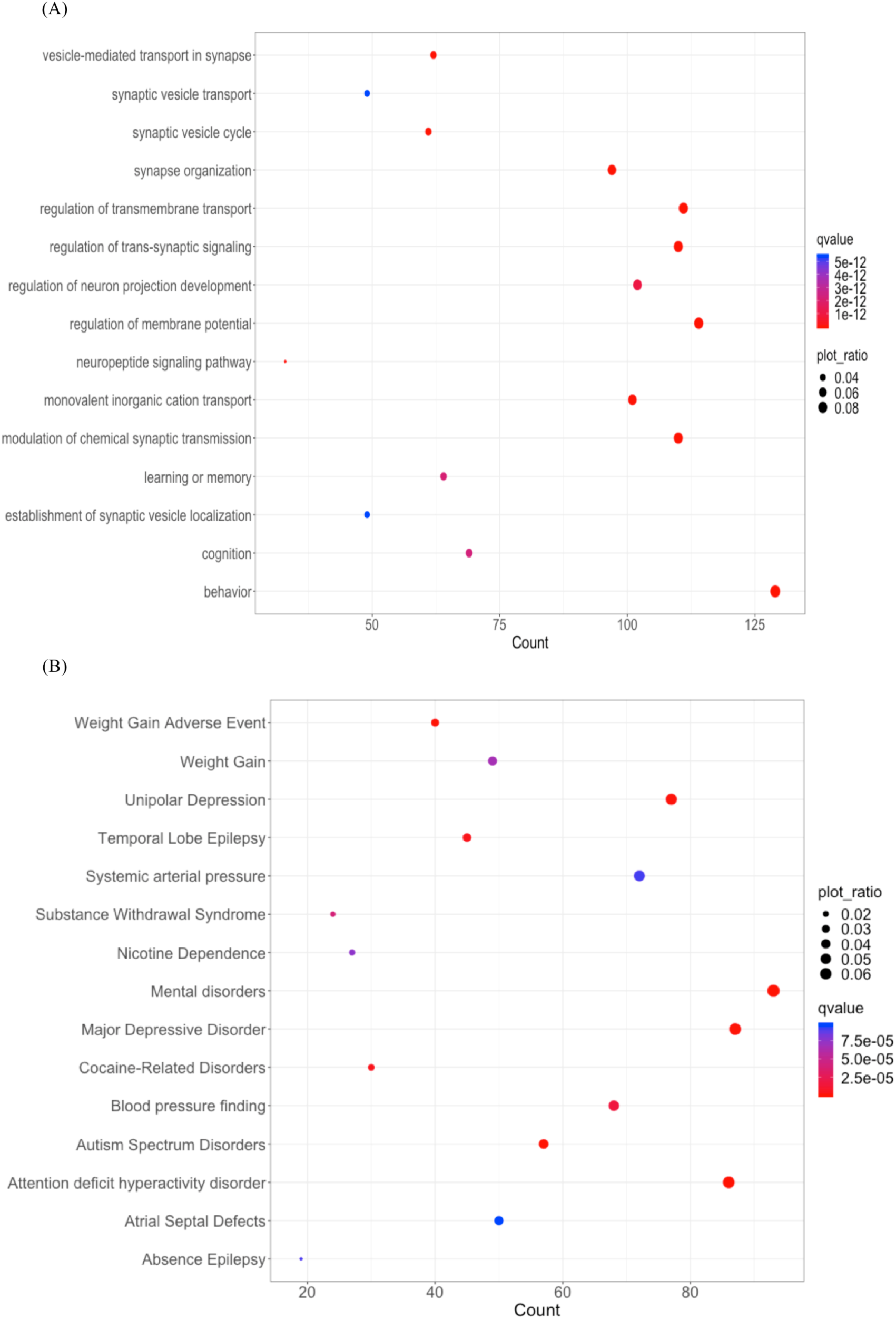
The functional enrichment of the parahippocampal gyrus brown module. The top 15 significant of (A) biological pathway (B) disease ontology enriched term of parahippocampal gyrus brown modules were plotted. The x-axis is the number of annotated genes and the y-axis is the enriched term. The dot size is proportional to the ratio of annotated genes/all background genes. The color of the dots represents significant qvalue.

**Supplementary Figure 5.**
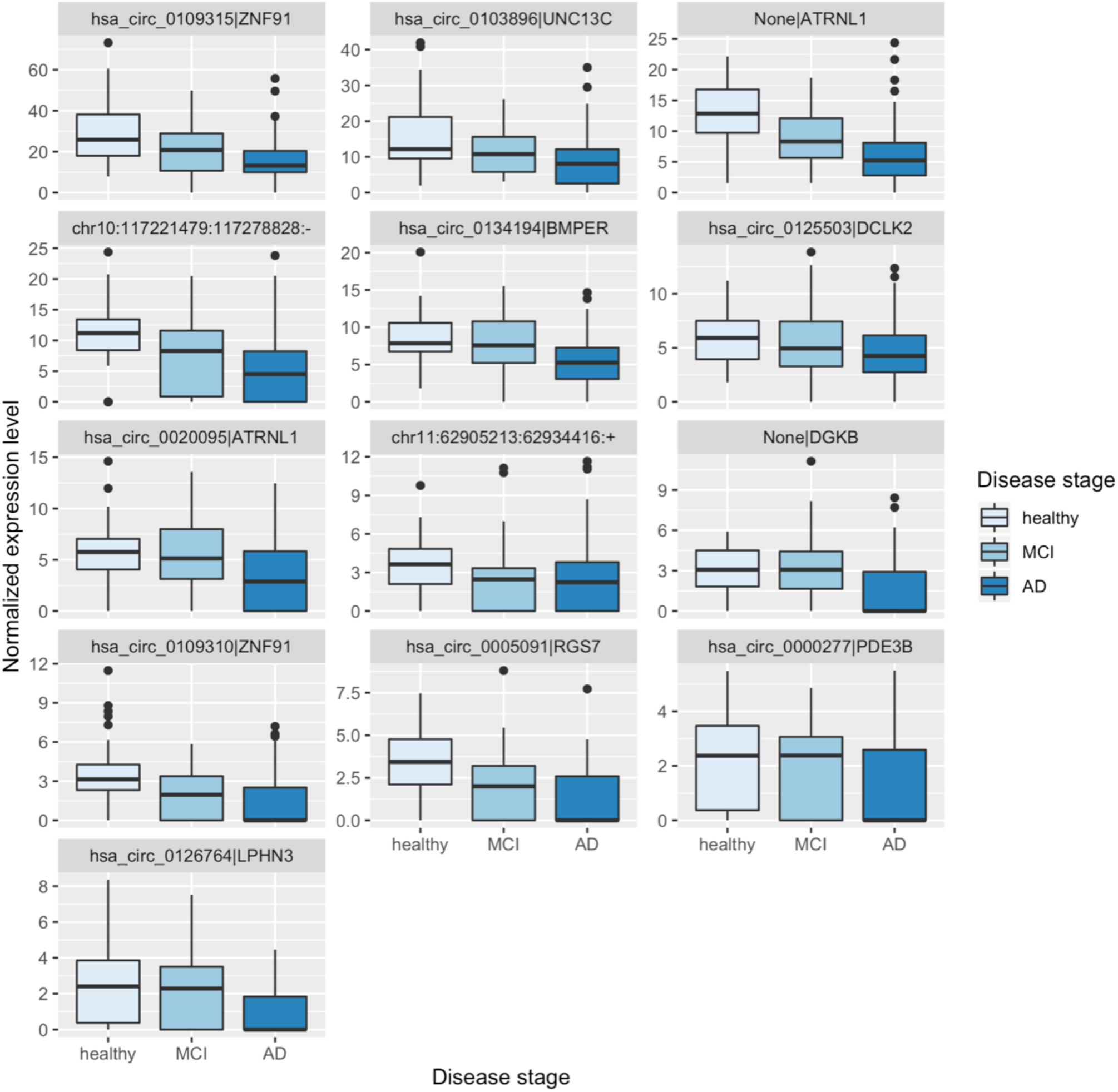
The circRNA expression level change of 13 selected circRNA candidates in different disease stage. The x-axis is the disease group and the y-axis is the expression level of each circRNAs.

**Supplementary Figure 6.**
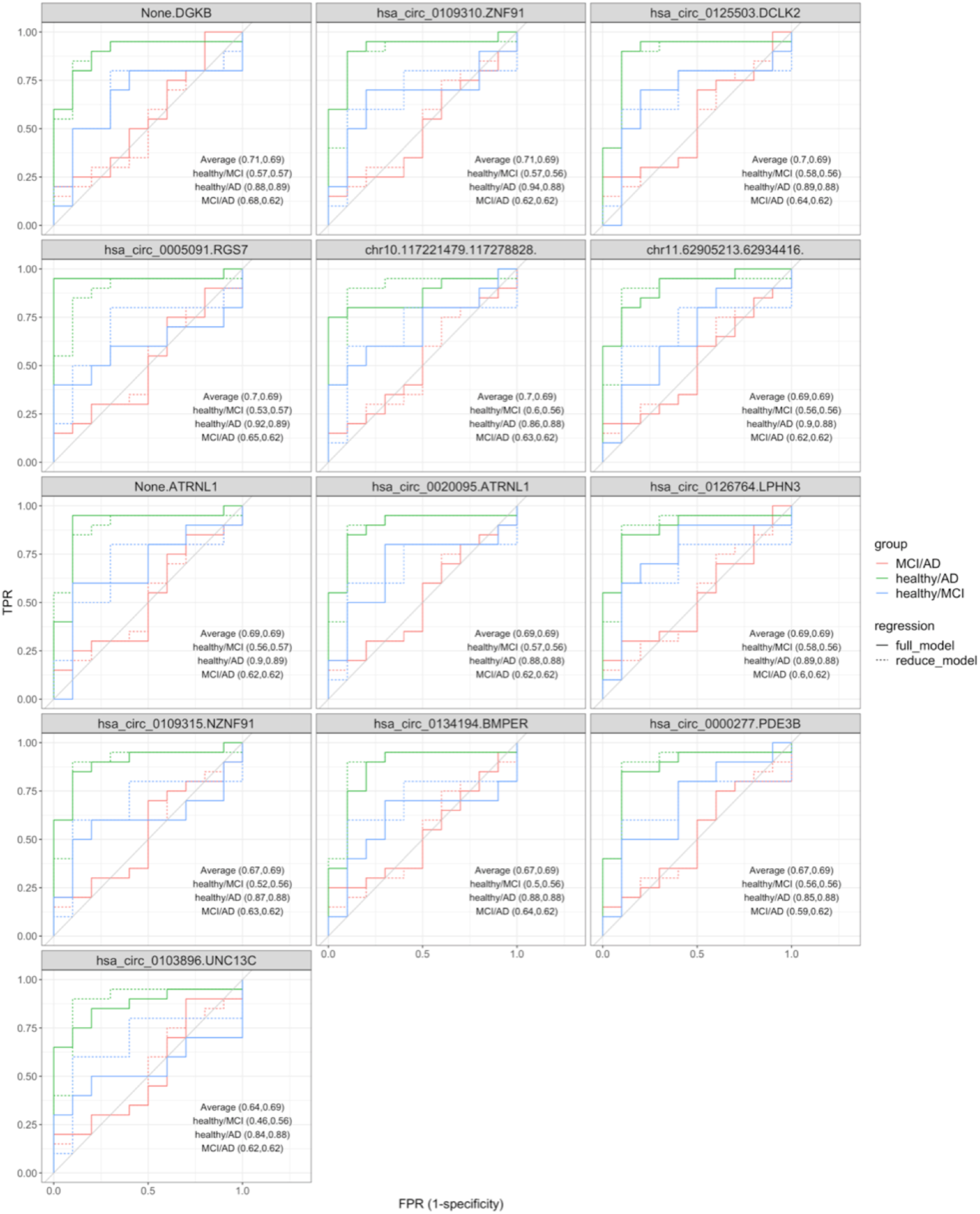
The ROC curve of multinomial logistic regression of 13 circRNA candidates. The solid line and dashed line represent the performance of the full and reduced model, respectively. The color shows the comparison between disease stages. The AUC of each ROC curve is annotated at the bottom right corner (the former number in the parentheses were AUC of the full model and the latter were AUC of the reduced model).

**Supplementary Figure 7.**
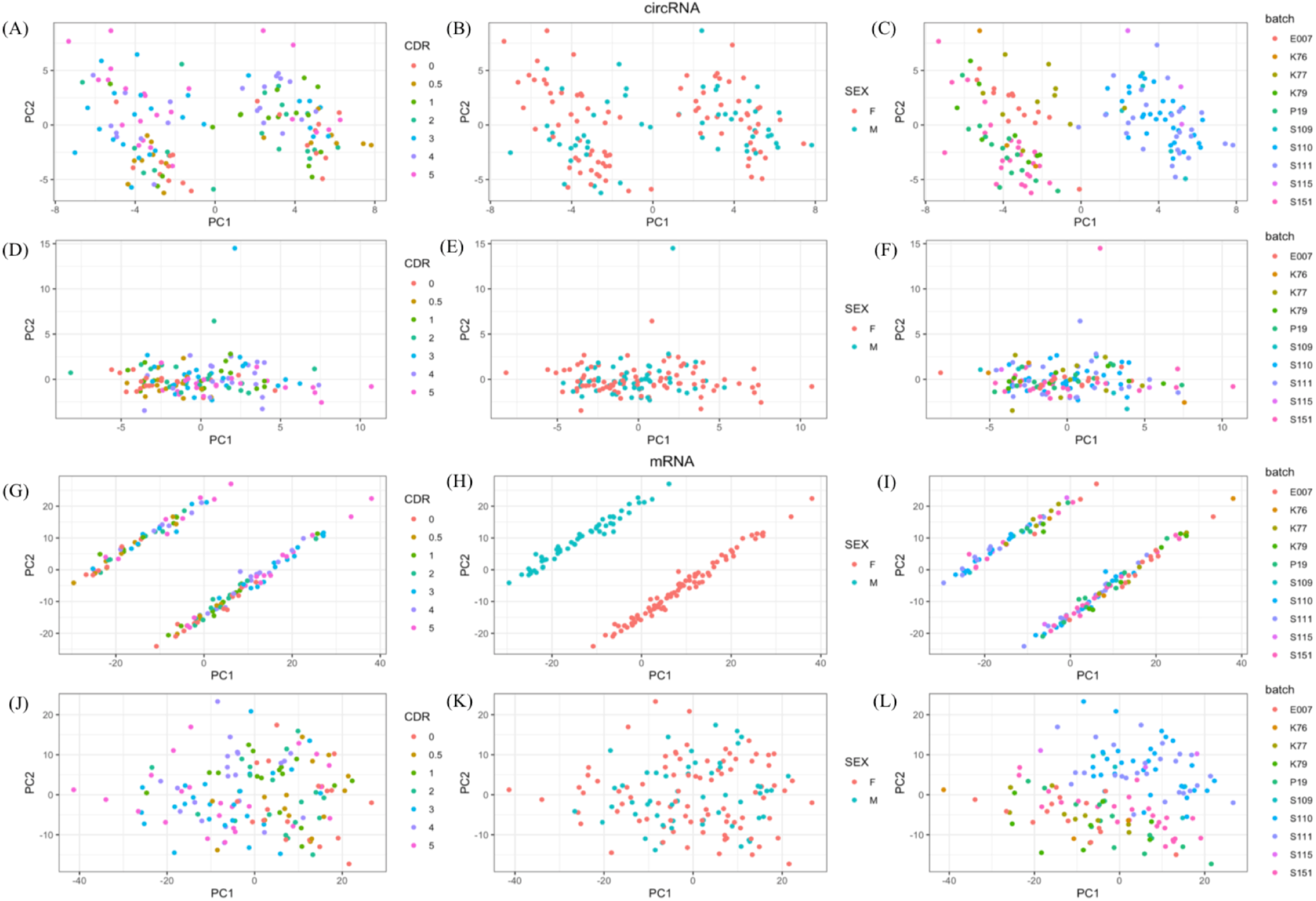
The confounder correction before co-expression network construction. (A)-(C) are the PCA of circRNA expression before limma correction and (D)-(F) are the PCA after limma correction, color code by CDR, sex, and batch, respectively. The PCA of mRNA expression before and after limma correction are shown in (G)-(I) and (J)-(L).

